# Stereotyped large amplitude cortical LFP events can be clustered and reveal precisely ordered phase-locking in neuronal populations

**DOI:** 10.1101/519454

**Authors:** Catalin C. Mitelut, Martin A. Spacek, Allen W. Chan, Tim H. Murphy, Nicholas V. Swindale

## Abstract

During quiet wakefulness, slow-wave sleep and anesthesia, mammalian cortex exhibits a synchronised state during which transient changes in the local field potential (LFP) accompany periods of increased single neuron firing, known as UP-states. While UP-state genesis is still debated (Crunelli and Hughes, 2010) such transitions may constitute the default activity pattern of the entire cortex (Neske, 2016). Recent findings of preserved firing order between UP-state transitions and stimulus processing in high-firing rate (>2Hz) rat auditory and barrel cortex neurons (Luczak et al., 2015) support this hypothesis. Yet it is unknown whether UP-states are homogeneous and whether neurons with firing rates <2Hz in visual cortex or other species exhibit spiking order. Using extracellular recordings during anesthetized states in cat visual cortex and mouse visual, auditory and barrel cortex, we show that UP-states can be tracked and clustered based on the shape of the LFP waveform. We show that LFP event clusters (LECs) have current-source-density profiles that are common across different recordings or animals and using simultaneous electrophysiology and widefield voltage and calcium imaging in mouse we confirm that LEC transitions are cortex-wide phenomena. Individual LEC events can be resolved in time to within 1 – 4 ms and they elicit synchronous firing of over 75% of recorded neurons with most neurons synchronizing their firing to within ±5 – 15 ms relative LECs. Firing order of different neurons during LEC events was preserved over periods of ~30 minutes enabling future studies of UP-state transitions and firing order with near millisecond precision.

**Significant Statement:** During sleep and anesthetic states mammalian cortex undergoes substantial changes from awake active states. Recent studies show that single neurons in some cortical areas in rats undergo increased spiking during sleep and anesthetic states (called UP-state transitions) with some neurons firing in an order similar to awake states. This suggests that sensory processing may be similar across all states and that firing order is important for stimulus processing. Yet UP-state transitions remain poorly understood and it is unclear whether firing order is present in other cortical areas or species. Here we describe multiple classes of UP-state transitions and show most neurons in visual cortex in cats and visual, barrel and auditory cortex in mice exhibit firing order during such transitions.

## Introduction

An established finding of the past several decades is that the cortical neurons of mammals spike differentially in two different states. One of these is a desynchronized state which is present during awake and attending periods and during rapid-eye-movement sleep (REM). During this state neurons fire largely independently of each other (Harris and Thiele, 2011). The other is a synchronized state, present during slow-wave sleep (SWS), quiet waking and anesthesia, where individual neurons cycle, at rates of 0.2 Hz – 0.9 Hz, between a depolarized (spiking) state and a hyperpolarized (non-spiking) state. These are known as UP- and DOWN-states, respectively (Steriade et al., 1993a; Sanchez-Vives and McCormick, 2000; McCormick and Yuste, 2006; Neske, 2016; Sanchez-Vives et al., 2017). UP- and DOWN-states are brain-wide phenomena which engage cortex, thalamus, hippocampus, striatum and cerebellum (Neske, 2016). They may facilitate flexible processing of information (McCormick et al., 2004; McCormick and Yuste, 2006; Haider et al., 2006) and may mediate changes in functional connectivity during waking states (Neske, 2016). They may also be involved in memory replay (Wilson et al., 1994; Sirota et al., 2003; Sirota and Buzsaki, 2005). While UP-state transitions can be evoked by sensory or thalamic activation (Amzica and Steriade, 1998; Steriade, 2001) transitions also occur spontaneously (Amzica and Steriade, 1995; Destexhe et al., 1999; Volgushev et al., 2006). Overall, slow oscillations and synchronized states might provide a “unifying paradigm for the study of cortical function” (Sanchez-Vives et al., 2017).

In addition to such broad scale functions the cortical “machinery” engaged by spontaneous UP-state transitions may be the same as used to represent stimuli during both awake, sleep or anesthetized states (Amzica and Steriade, 1998). A number of extracellular recording studies found that high firing rate (>2Hz) neurons in rat somatosensory and auditory cortex tend to fire in a similar order during UP-state transition (though some firing distributions were broad, e.g. 50-100ms full-width-half-max) as well as during the first 100ms following stimulus onset (Luczak et al., 2007, 2009; Bermudez-Contreras et al., 2013; Luczak et al., 2015). Firing order during UP-state transitions may therefore reveal a functional role for order ni cortex, and tracking spiking order for all cells and in other cortical areas might help reveal a coding strategy used by cortex. While traditional methods for defining UP-state transitions rely on single-cell intracellular recordings (Steriade et al., 1993a), these methods can only be used to track UP-states for a few neurons at a time. Additional methods for UP-state detection also have relied on peaks in synchronous firing (Luczak et al., 2007), but such methods exclude low-firing rate neurons (e.g. neurons firing <2Hz) and sparsely firing cortical areas (e.g. visual cortex).

Here we expand the analysis of UP-states by providing a method for detecting them using only extracellular recordings and show that most neurons (even those with low firing rates) have a preserved firing order. We show that the multi-channel local field potential (LFP) generated during UP-state transitions can be clustered and provides a more temporally precise definition of UP-states. Previous studies have shown that single-channel LFP events correlate with UP-state transitions (Saleem et al., 2010; Chauvette et al., 2010) and more recent work in rat hippocampal slices (Reichinnek et al., 2010) and anesthetized macaque hippocampus (Ramirez-Villegas et al., 2015) have shown that LFP events can have stereotyped shapes. Here we go further and show that in the synchronized state, large amplitude channel LFP events can be clustered (termed LFP event clusters - LECs) within a 1 – 4ms temporal precision enabling the accurate measurement of latencies of simultaneously recorded single units. CSD analyses of LECs revealed characteristic laminar profiles of sources and sinks for each LEC and potentially 3 groups of such clusters. We additionally used widefield voltage-sensitive dye (VSD) imaging of mouse cerebral cortex to show that LECs are associated with cortex-wide activations consistent with previous work (Amzica and Steriade, 1998). Most neurons synchronized their firing to individual events to within ±5 – 15 ms. Consistent with this, the relative firing order of different units during LFP events was present in cat visual cortex and mouse visual, barrel and auditory cortex adding to the evidence that cortical neurons are capable of firing with high temporal precision relative to each other.

## Materials and Methods

### Experimental Design

#### Cat Electrophysiological Recordings

Experimental procedures are described in detail in previous work (Swindale and Spacek, 2014) and were carried out in accordance with guidelines established by the Canadian Council on Animal Care and the Animal Care Committee of the University of British Columbia. Data analysed here were obtained from 15 electrode penetration sites in 5 adult cats (animal IDs: C1-C5). The cats were anesthetized either with 0.5-1.5% isoflurane and 70% N2O + 30% O2 (C1, C2 and C3) or with continuously infused propofol (6 – 9 mg/kg/hr) and fentanyl (4–6 *μ*g/kg/hr) (C4, C5). Following craniotomy surgery, a high-density polytrode was inserted perpendicularly into the cortex until the upper recording sites were 100 – 200 *μ*m below the surface. Polytrodes were either 2-column (C2,C4,C5) or 3-column (C1, C3) with electrode site spacing of 50-75 *μ*m. Voltage signals were analogue bandpass filtered between 0.5 and 6 kHz, sampled at a rate of 25 kHz and digitized with 12-bit resolution (Blanche et al., 2005). A subset of 10 electrode sites were used to separately record the LFP, band pass filtered between 0.5-200 Hz, were fed in parallel to separate amplifiers. On the 3-column electrodes the channels were 130 *μ*m apart, with the exception of the bottom 2 channels, which were 65 μm (C1) or 97 *μ*m (C3) apart. On the 2-column electrodes the channels were 150 *μ*m (C2 and C4) or 195 *μ*m (C5) apart, with the lower two channels being 100 *μ*m (C2 and C4) or 195 *μ*m (C5) apart. Recording sites were in area 17 and receptive fields (not reported here) were typically within 10 degrees of the *area centralis*. In addition to recordings of spontaneous activity, visual stimuli, including moving bars, gratings, m-sequence stimuli and natural scene movies were presented on a CRT screen. Table 1 summarizes recording IDs, anesthetic methods and recording duration for individual experiments in cats.

**Table 1:** Cat Visual Cortex – Recording Summary

| Rec ID | Anesthetic | Track ID | Synchronized Rec Time<sup>1</sup> /Total Rec Time | No. of LECs |
| --- | --- | --- | --- | --- |
| C1.1 | Iso/N20 | 1 | 0.6 / 13.4hrs<sup>2</sup> | 2 |
| C1.2 | Iso/N20 | 2 | 1.4 / 9.4hrs<sup>2</sup> | 2 |
| C2.1 | Iso/N20 | 1 | 1.7 / 11.1hrs<sup>2</sup> | 3 |
| C2.2 | Iso/N20 | 2 | 4.7 / 12.0hrs<sup>2</sup> | 1 |
| C3.1 | Iso/N20 | 1 | 1.4 / 8.4hrs | 1 |
| C3.2 | Iso/N20 | 2 | 4.2 / 8.3hrs | 3 |
| C3.3 | Iso/N20 | 3 | 3.4 / 7.4hrs | 2 |
| C3.4 | Iso/N20 | 4 | 3.0 / 5.3hrs | 1 |
| C4.1 | Prop/Fent | 1 | 3.3 / 11.9hrs | 1 |
| C4.2 | Prop/Fent | 2 | 0.5 / 11.5hrs | 2<sup>3</sup> |
| C4.3 | Prop/Fent | 3 | 0.9 / 4.6hrs | 2 |
| C5.1 | Prop/Fent | 1 | 2.5 / 2.9hrs | 3 |
| C5.2 | Prop/Fent | 2 | 4.9 / 10.6hrs | 3 |
| C5.3 | Prop/Fent | 3 | 3.1 / 8.5hrs | 4 |
| C5.4 | Prop/Fent | 4 | 4.6 / 6.3hrs<sup>2</sup> | 4 |
| Totals |  | 15 |  | 34 |

1 – Synchronized recording time was defined as periods with synchrony index > 0.5.
2 – Synchrony index of 0.3 used.
3 – LECs had peak-to-peak amplitude and CSD maxima below threshold and were excluded from CSD grouping and further analysis.

#### Mouse Electrophysiological Recordings

Experimental protocols were established and carried out in accordance with guidelines established by the Canadian Council on Animal Care and the Animal Care Committee of the University of British Columbia. Data reported here were obtained from a total of 4 electrode tracks in 4 mice (C57/BL6) anesthetized with isoflurane (1.5–2%) for surgery and with subsequent recording periods under reduced concentration of isoflurane (1.0–1.2%). The skull was fixed to a head-plate to stabilize recording and facilitate imaging in simultaneous acquisition sessions. Extracellular recordings were made with 64-channel polytrodes (NeuroNexus A1×64-Poly2-6mm-23s-160-A64) with a 2-column (32 channels per column) staggered-format with vertical and horizontal (inter-column-distance) of 46 μm covering 1450 μm of the probe. Voltage signals were acquired using a headstage amplifier (RHD2164, IntanTech, Los Angeles) and USB interface board (RHD2000, Intan) at a sampling rate of 25 kHz - 16 bit resolution. Electrodes were inserted perpendicular to the surface of the cortex using a micro-manipulator (MP-225, Sutter Instrument Company). Cortical penetration depth was tracked using micro-manipulator coordinates with the tip of the electrode being inserted between 900 μm to 1450 μm (mean of 1256 μm ± 157 μm) below the cortical surface. Further details for these recordings can be found in (Xiao et al., 2017).

#### Mouse VSD Imaging

To determine the cortex-wide correlates of LECs, widefield VSD imaging was carried out in anesthetized mice as previously described (Mohajerani et al., 2010, 2013; Vanni and Murphy, 2014) while simultaneously recording LFP and single unit activity extracellularly. Either a unilateral craniotomy (1 wildtype C57/BL6 mouse, from bregma 2.5 to −4.5 mm anterior-posterior, and 0 to 6 mm lateral) or a bilateral craniotomy (2 wildtype C57/BL6 mice, from bregma 3.5 to −5.5 mm anterior-posterior, and −4.5 to 4.5 mm lateral) was made and the underlying dura removed. RH1692 dye (Optical Imaging, New York, NY; (Shoham et al., 1999) dissolved in HEPES-buffered saline (1 mg/ml) was added to cortex for 60–90 min. VSD imaging began ~30 minutes following washing of unbound dye with saline. VSD data (12 bit monochrome) was captured with 6.67 ms (150Hz) temporal resolution using a CCD camera (1M60 Pantera, Dalsa, Waterloo, ON) and EPIX E4DB frame grabber with XCAP 3.1 software (EPIX, Inc., Buffalo Grove IL).

Table 2 summarises recording IDs, anesthetic methods and recording duration for individual extracellular and VSD recording experiments in mice.

**Table 2:** Mouse Cortex – Recording Summary

| Rec ID | Anesthetic | Track ID | Area | Imaging | Sync Rec Time /Total Rec Time | No. of LECs |
| --- | --- | --- | --- | --- | --- | --- |
| MV1 | Isoflurane | 1 | Visual | VSD | 2.6 / 3.1hrs | 2 |
| MV2 | Isoflurane | 1 | Visual | VSD | 2.2 / 2.2 hrs | 1 |
| MV3 | Isoflurane | 1 | Visual | No | 2.5 / 2.5 hrs | 2 |
| MV4 | Isoflurane | 1 | Visual | GCaMP6s | 1.0 / 1.0 | 1 |
| MA1 | Isoflurane | 1 | Auditory | VSD | 2.1 / 2.1 hrs | 1 |
| MA2 | Isoflurane | 1 | Auditory | GCaMP6s | 3.1 / 3.1 hrs | 1 |
| MA3 | Isoflurane | 1 | Auditory | No | 2.0 / 2.0 hrs | 1 |
| MB1 | Isoflurane | 1 | Barrel | GCaMP6s | 2.7 / 2.7 hrs | 1 |
| Totals |  | 8 |  |  |  | 9 |

### Analysis

Most of the analyses were carried out using custom Python code developed as part of an electrophysiology and optical physiology toolkit currently in development (https://github.com/catubc/openneuron). Methods for computing event triggered analysis for VSD imaging have been previously published (Xiao et al., 2017) and are also available online (https://github.com/catubc/sta_maps).

#### Single Unit Spike Sorting

Spike sorting of cat and mouse data was carried out primarily using SpikeSorter (Swindale and Spacek, 2014, 2015) and selectively using Kilosort (Pachitariu et al., 2017) and JRClust (Jun et al., 2017). For recordings sorted using SpikeSorter, electrophysiological traces were high-pass filtered and spikes detected using a threshold of 5 times the median of the absolute voltage values of each channel was divided by 0.675 (Quian Quiroga et al., 2004) followed by a dynamic-multiphasic event detection method (Swindale and Spacek, 2015). A summary of the sorting results is provided in Table 3 and examples of sorted spike waveforms are shown in Supplementary Figure S3.

**Table 3:**
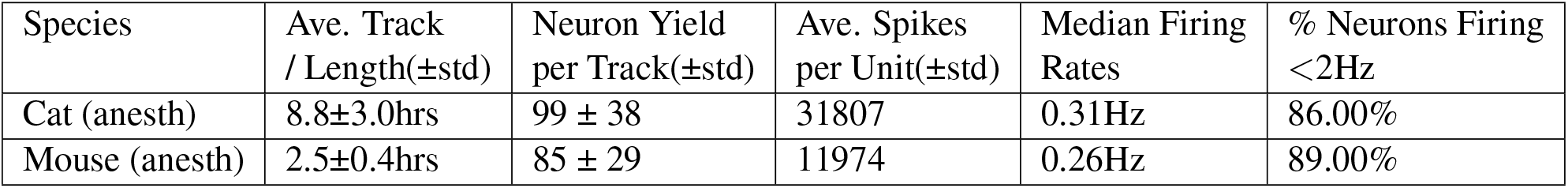
Single Unit Sorting Summary

#### Clustering LFP Events

High-amplitude LFP events were detected and clustered by converting LFP recordings to a data format similar to that of a high-pass spike recording. Existing spike sorting tools were then used for event detection, alignment, feature-extraction, clustering and review. Synchronized states (Figs 1A) were identified using the deepest LFP channel and on the basis of a synchrony index (SI) (Li et al., 2009; Saleem et al., 2010) which measures the ratio between power below 4 Hz and total power. Values of SI greater than 0.5 (i.e. periods where most of LFP power lies in the 0.1 – 4 Hz band) were used to define the synchronous state. In cat V1 recordings, synchronized state periods accounted for 2.7 ± 1.5 hrs out of a total recording time of 8.8 ± 3.0 hrs (Table 1) but varied substantially for each recording ranging from 4% to 86% of the total recording period for each animal. This was likely due to variability of anesthetic depth and animal physiology. In anesthetized mouse sensory cortex recordings, synchronized state periods accounted for 2.3 ± 0.2 hrs ranging from 84% to 100% of the total recording periods (Table 2).

**Figure 1.**
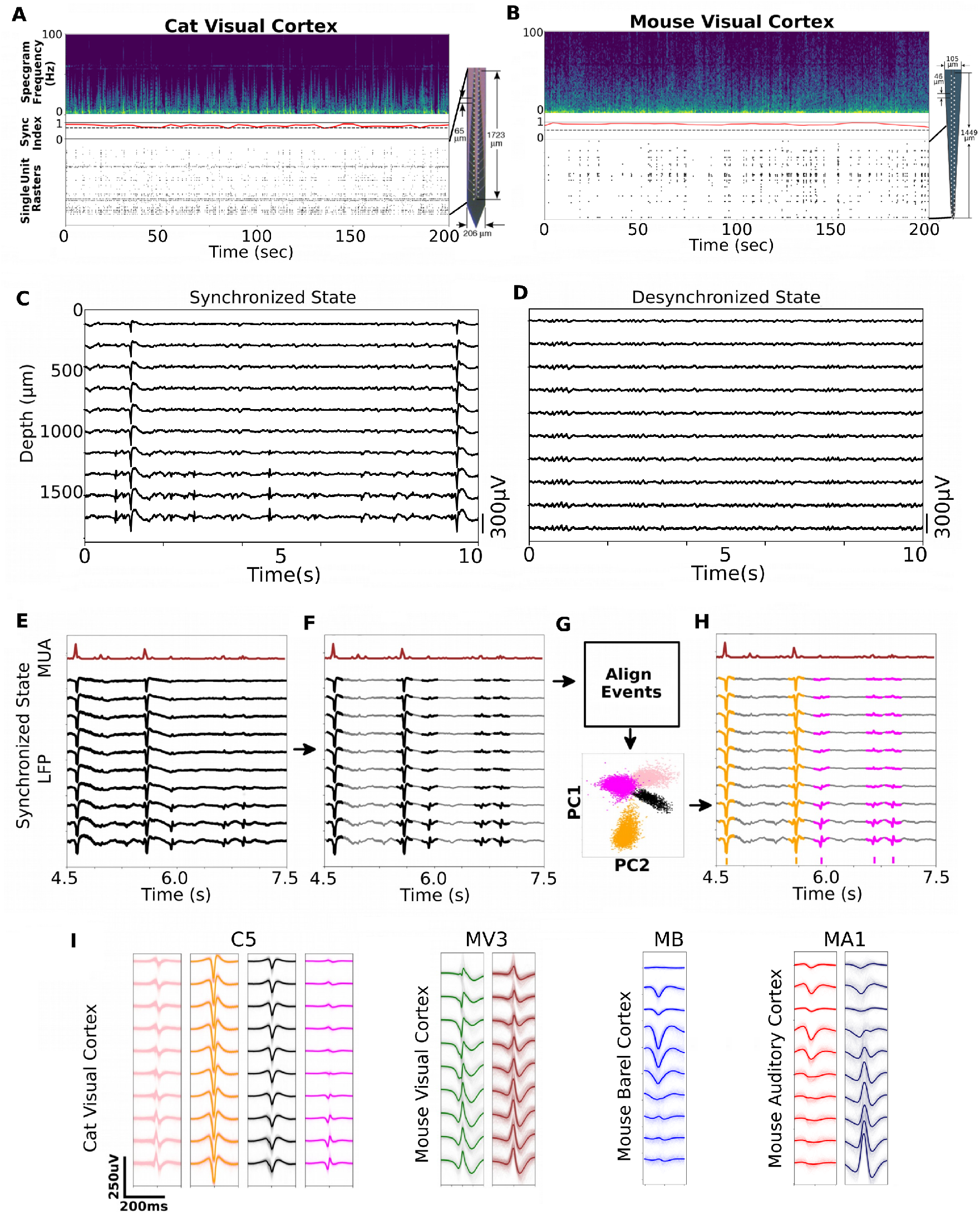
Clustering LFP Events During Synchronized Cortical States. A. Example of a 200 s cat visual cortex extracellular recording with power spectrogram (top), synchrony index (middle), single neuron rasters (bottom) and 64 channel extracellular probe diagram (right). B. Same as (A) but from mouse visual cortex using a different type of extracellular probe containing 64 channels. C. Extracellular recording of 10-LFP channels obtained from a track in cat visual cortex during a synchronized state reveals infrequent large amplitude cross-laminar events (animal ID: C5). D. Same track as in (A) but recording acquired during a desynchronized cortical state shows higher frequency cross-laminar events that are much smaller. E. Large amplitude LFP events correlate with peaks in MUA histograms during synchronized state recordings (ID: C5). F. Large amplitude LFP events are detected (see text for details). G. detected events are aligned and features are clustered using PCA. H. LFP events are labeled (colors) and event times exported for analysis (see also Methods). I: left four panels: LFP templates for the 4 LECs identified in (F) show distinct multi-laminar LFP patterns. Middle and right panels: LFP templates for LECs identified in mouse visual, barrel and auditory cortex respectively.

Synchronized state LFP recordings were next high-pass filtered with a 4-pole Butterworth filter with a cutoff of 4 Hz to remove slower LFP fluctuations and improve signal-to-noise ratio (SNR) for subsequent event detection, alignment and clustering. Next, event-detection was performed using the same methods described above for spikes, with the same detection threshold and a temporal window of 50 ms (Swindale and Spacek, 2015). A temporal lockout of 150 ms and a spatial lockout of 2 mm were used to ensure that in a 150 ms period only a single LFP event could be identified. Following detection, events were initially aligned using a weighted center of gravity definition of the time of the event (Swindale and Spacek, 2014). Max peak alignment revealed similar results. Principal components were then calculated based on the covariance matrix obtained from the 50 points with the highest voltage variance taken across all channels. The observed principal component value distributions normally showed clear evidence of clustering (Figs. 1E, 2). Clustering was done based on the first two principal component values. Following clustering, the mean waveform of the events in the cluster was calculated and the individual waveforms were then further aligned to this mean using r.m.s. error minimization (Swindale and Spacek, 2014) computed over all the LFP channels. The mean waveform was then recalculated and the process was repeated until further realignments were of vanishingly small magnitude. The time in the aligned event waveform that corresponded to the center of gravity of the template (defined as above for individual waveforms) was then taken as the time of the event. Note that any other stable feature of the template could equally well have been used as an anchor as this would simply change the times of all the events by the same amount. We used the center of gravity measure in preference to peaks, troughs or zero-crossings as these features can be variable across different LECs and are occasionally ambiguous.

**Figure 2.**
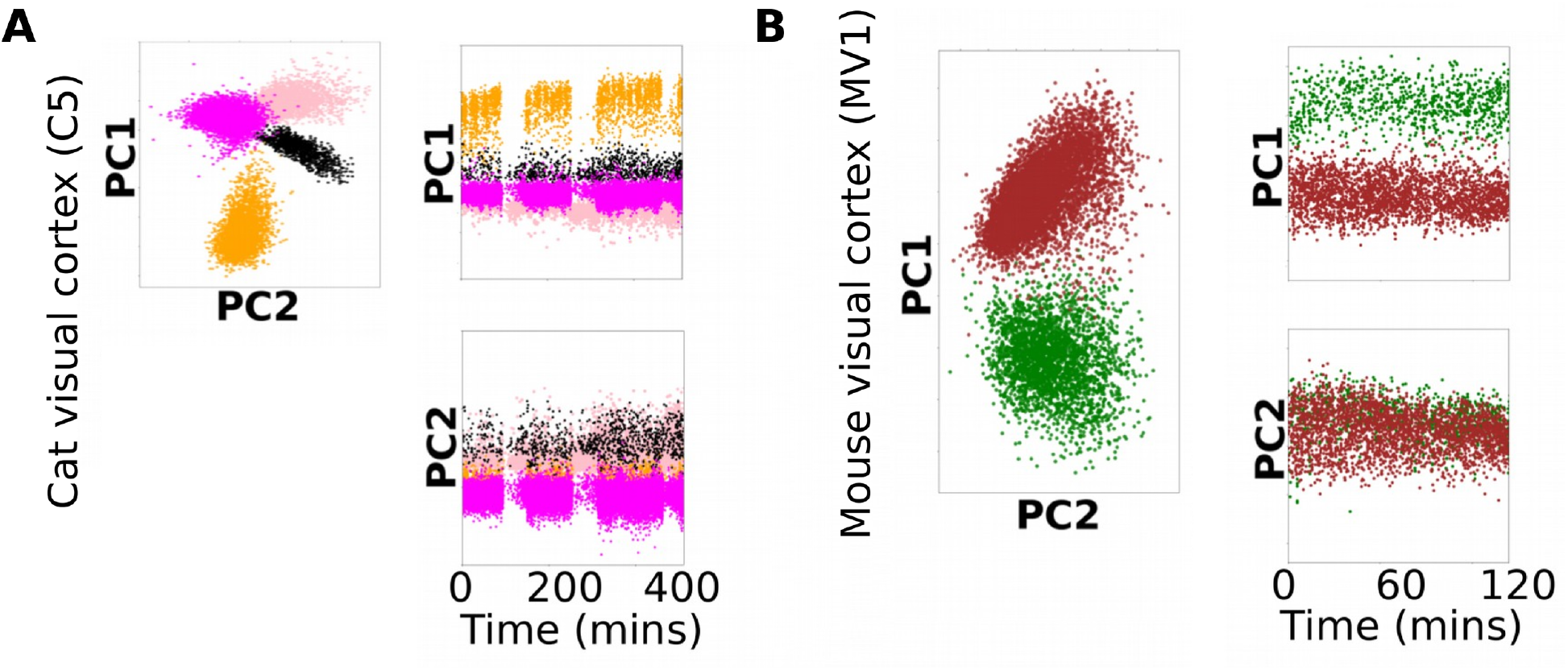
Stability of LECs over Time. A: shows the principal component values (PC1 and PC2) for the 4 LECs shown in Fig 1E, plotted over a period of 400 minutes (6.7 hours). This shows that individual LEC features are relatively stable over time. B: same as A but for 2 LECs in a mouse visual cortex recording.

Clusters with < 20 events or with peak-to-peak heights of < 100 uV were deleted and excluded from further analysis.

#### Precision of Estimation of Event Times

The accuracy with which individual LEC events can be aligned to the template in the presence of background variability in the LFP signal determines the accuracy of the estimate of the times at which individual events can be said to have occurred. This accuracy potentially limits the ability to determine the variability in the timing of spikes of individual units relative to the event. If the accuracy is low, the measured variability in timing will be larger than it actually is. We estimated the accuracy of the r.m.s. alignments by computing hybrid ground truth data (i.e. using real data with simulated shifts). We first calculated the covariance matrix of the noise in the LFP signal relative to a particular LEC template. This was normally based on the noise values for 100 points (1 ms) on the LFP channel for which the LEC peak-to-peak amplitude was a maximum. Alternatively, several different channels were included in the noise calculation. This matrix was then used to generate simulated noise samples with the same amplitude and covariance structure as the real noise. Simulated LEC waveforms were then generated by adding samples of simulated noise to the LEC template. Each waveform was then realigned to the template using r.m.s. minimisation and the resulting shift in position was taken as the error for that particular sample. The mean of the errors was normally close to zero and the standard deviation of the errors measured for 1000 random samples was taken as an estimate of the accuracy.

#### CSD Computation

LEC CSDs were computed by calculating the second spatial derivative (Nicholson and Freeman, 1975) of LEC templates using all available LFP channels. This calculation was implemented using the gradient function of the numpy Python library which provides “first or second order accurate one-sides (forward or backwards) differences at the boundaries” (https://docs.scipy.org/doc/numpy-1.13.0/reference/generated/numpy.g

#### LEC-Triggered VSD Motifs

VSD motifs were computed as described by (Xiao et al., 2017). A response, dF/F0, was computed for −3 s to +3 s around each LEC event, with F0 calculated as the average of the signal −6s to −3s before each event. Strong sensory stimulation resulted in VSD signals which generally peaked at 0.5% dF/F0. The LEC triggered VSD motifs had peaks of 0.1-0.2%. These were substantially larger than randomly generated motif peaks (See Fig 8A-control).

#### Grouping of LECs using CSD shapes

CSDs were equalized and then clustered using a generalized mixture model with 3 components (Fig 4). Because the recordings were made with different length electrodes, all CSDs shapes were clipped to represent only 0 *μ*m to 1200 *μ*m of cortical tissue. This allowed for a proper comparison to be made across all 24 selected recordings. Next, the 2D-shape CSDs were aligned to the mean of all 2D shapes and then converted to a 1D vector. The 1D vector array for all CSDs was then compressed using principal component analysis and a generalized mixture model with n=3 components was fit to the resulting distributions (3 was chosen as qualitatively there appeared to be 3 different shapes considering both CSD shapes and PCA distributions).

## Results

### Clustering Large Amplitude Multi-Laminar LFP Events Reveals Distinct Event Classes

Multi-channel extracellular recordings were made during synchronized states in anesthetized cat visual cortex and mouse visual, barrel and auditory cortex (Fig 1A, B). Synchronized state recordings in anesthetized cats and mice contained large-amplitude, stereotypically shaped LFP events (Fig. 1C). These large amplitude events were mostly absent during desynchronized cortical states where only lower amplitude events were typically observed (Fig. 1D). Large amplitude LFP events correlated with peaks in multi-unit-activity (Fig 1E) which are commonly known to be global indicators of UP-state transitions (e.g. (Luczak et al., 2007)). The LFP events generally had 1 – 3 peaks and troughs with varying relative heights and widths (Fig. 1F). Peak-to-peak amplitudes were typically 250 – 500 μV (after high-pass filtering at 4 Hz) and the duration of the events was typically 50 – 100 ms. Clustering these events (see Methods) we identified 1 - 4 LECs per recording session in cat visual cortex and 1 - 2 LECs in mouse visual, auditory and barrel cortex recording sessions (Fig 1H, I; Tables 1 and 2; see also Methods).

The stability of LEC shapes could be tracked over time using the principal component values of the LEC event waveforms. We found that LEC shapes were stable and the principal component values of events within each cluster did not change substantially over periods of up to 3 hours in either cat or mouse cortical recordings (Fig. 2A, B).

### LECs Reflect Distinct CSD Patterns Common Within and Across Animals

We used CSD analysis to further investigate the properties of LECs within and across animals (Figs. 3 and 4). Both the raw unaveraged LFP traces (Fig. 3A, B, E, G) and the averaged LEC profile (Fig. 3C, D, F, H) revealed distinct CSD patterns present in cortex during synchronized states. CSD profiles from synchronized state cortical recordings showed periodic large amplitude events corresponding to LECs (Fig 3A, E and G) whereas CSD profiles from desynchronized state recordings contained more frequent activity distributed across multiple depths and with lower amplitude (Fig 3B; note amplitude were normalized within each session). Computing CSD profiles for different LEC templates in cat V1 recordings current-sink distributions showed they could be similar across different tracks within the same animal. For example, Fig. 3C shows 4 similarly shaped LEC CSD profiles from 4 different tracks in two hemispheres from a single cat. Some profiles were similar across different animals. Fig. 3D shows 5 similarly shaped CSDs, 4 of which are from four tracks in one cat in both hemispheres and 1 from another cat. Figures 3E and G show CSD profiles for unaveraged LFP events recorded in mouse visual and auditory cortex respectively. Figures 3F and H show CSD profiles obtained from the averaged LEC waveform for mouse visual and auditory cortex respectively.

**Figure 3.**
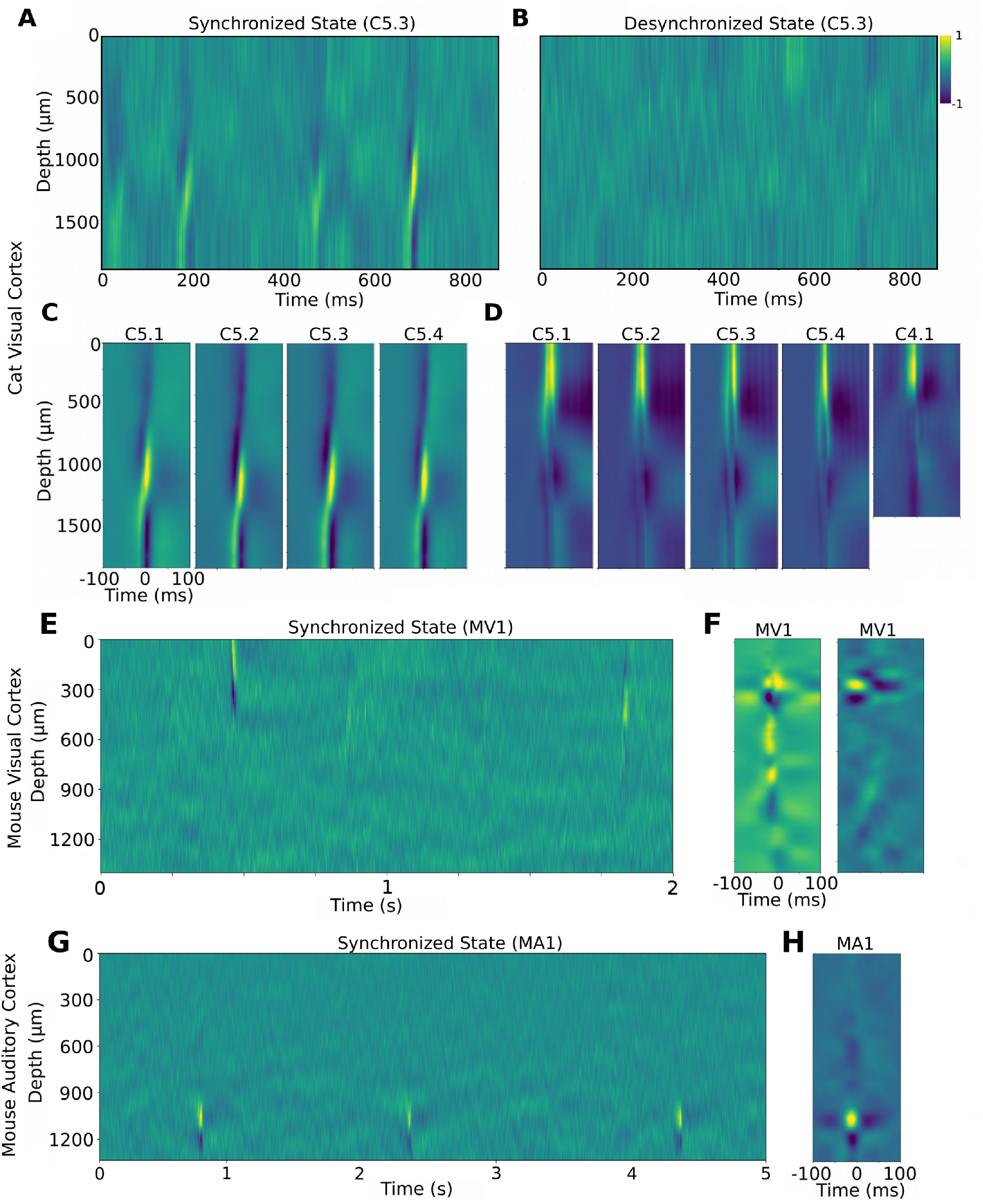
Current-Source-Density (CSD) Analysis of LECs. A. CSDs for a 850 ms synchronized state recording (same track in animal C5 as shown in Fig 1A). B. CSD distributions for a 850 ms desynchronized state recording (same track as Fig 1B, 3A). C. CSD obtained from averaged LEC event profiles (i.e. templates) recorded in 4 different visual cortex tracks in animal C5. D. CSD distributions for averaged LEC profiles recorded in 5 different visual cortex tracks in two cats (1-4: animal C5; 5: animal C4). E. Same as panel A, but for a recording from mouse visual cortex. F. CSDs from two averaged LEC waveforms in a single mouse visual cortex recording. G. CSD profiles from unaveraged LFP events in a synchronised recording from mouse auditory cortex. H. Same as 3C, F but for a mouse auditory cortex recording. All CSD distributions were normalized to lie between −1 and +1 for visualization.

**Figure 4.**
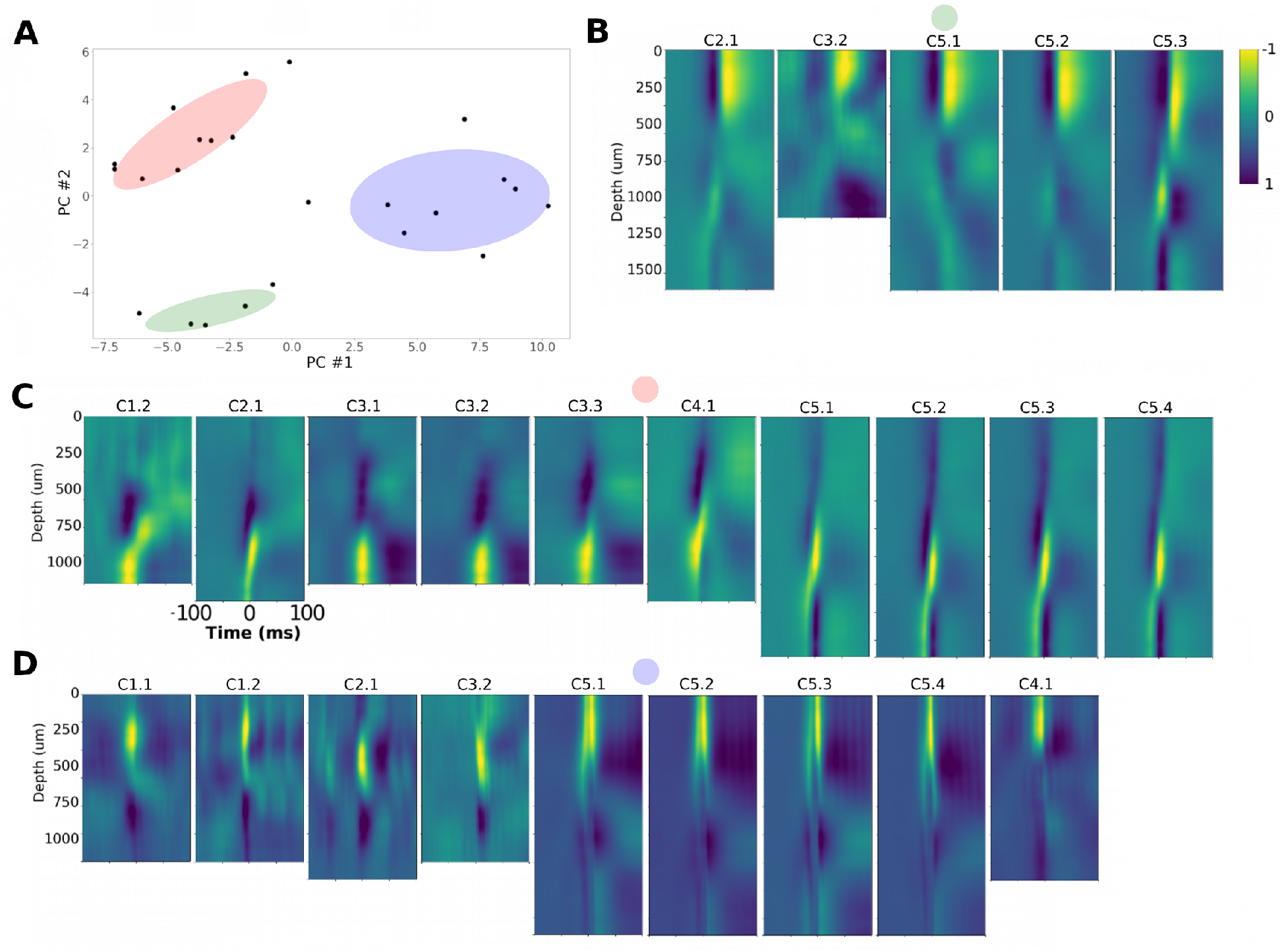
LEC Grouping and LEC Shape Stability. A. Principal component distributions for the most common 24 CSDs detected in cat visual cortex recordings were fit with a 3-component generalized gaussian mixture model (see Methods). Shaded regions represent 1 standard deviation. B-D. CSD for each group clustered in A.

### LECs Are Similar Within and Across Animals

Across 14 V1 cortex tracks in 5 cats we identified 34 LECs (Table 1) with between 1 – 4 LEC per track. Using a gaussian mixture model with 3 components we grouped the 24 most common CSD shapes into three groups (Fig 4; see Methods). Thus rather than being unique or specific to a particular animal or track, LECs can be grouped on the basis of their CSD shape and can be the same within and across different cats. This suggests LECs may be generated by common neural circuits underlying UP-state transitions. Different types of UP-state transitions may thus be present across animals within a species (Table 4). Additionally, in a total of 8 mice recordings from visual, auditory and barrel cortex we found 13 LECs that could also be grouped (Table 7). The average frequency of the LECs (taken across all types) was 0.12 Hz across all cat V1 recordings, and 0.18 Hz across all mouse cortex recordings.

**Table 4:** Cat V1 – LEC Groupings Summary

| Rec ID | Anesthetic | #1 | #2 | #3 | #4 | #5 | #6 | Other |
| --- | --- | --- | --- | --- | --- | --- | --- | --- |
| C1.1 | IsoN20 |  | X |  |  |  |  | X |
| C1.2 | IsoN20 | X | X |  |  |  |  |  |
| C2.1 | IsoN20 | X | X |  |  |  |  |  |
| C2.2 | IsoN20 |  |  |  |  |  | X |  |
| C3.1 | IsoN20 | X |  |  |  |  |  |  |
| C3.2 | IsoN20 | X | X | X |  |  |  |  |
| C3.3 | IsoN20 | X |  |  |  | X |  |  |
| C3.4 | IsoN20 |  |  |  | X |  |  |  |
| C4.1 | Prop/Fent | X | X |  |  |  |  |  |
| C4.2 | Prop/Fent |  |  |  |  |  | X | X |
| C5.1 | Prop/Fent | X | X | X |  |  |  |  |
| C5.2 | Prop/Fent | X | X | X |  |  |  |  |
| C5.3 | Prop/Fent | X | X | X |  |  |  | X |
| C5.4 | Prop/Fent | X | X | X |  |  |  |  |
| Totals |  | 10 | 9 | 5 | 1 | 1 | 2 | 3 |

1 – These 2 LECs are similar to Class 6 but may form separate class.

**Table 5:** Mouse Cortex – LEC Groupings Summary

| Rec ID | Anesthetic | Area | #1 | #2 | #3 | #4 | #5 | #6 | #7 | #8 |
| --- | --- | --- | --- | --- | --- | --- | --- | --- | --- | --- |
| MV1 | Isoflurane | Visual | X | X |  |  |  |  |  |  |
| MV2 | Isoflurane | Visual | X |  |  |  |  |  |  |  |
| MV3 | Isoflurane | Visual | X |  | X | X |  |  |  |  |
| MV4 | Isoflurane | Visual | X |  |  | X |  |  |  |  |
| MA1 | Isoflurane | Auditory |  |  |  |  | X | X |  |  |
| MA2 | Isoflurane | Auditory |  |  |  |  |  | X |  |  |
| MA3 | Isoflurane | Auditory |  |  |  |  |  |  | X |  |
| MB1 | Isoflurane | Barrel |  |  |  |  |  |  |  | X |
| Totals | 8 |  | 4 | 1 | 1 | 2 | 1 | 2 | 1 | 1 |

**Table 6:**
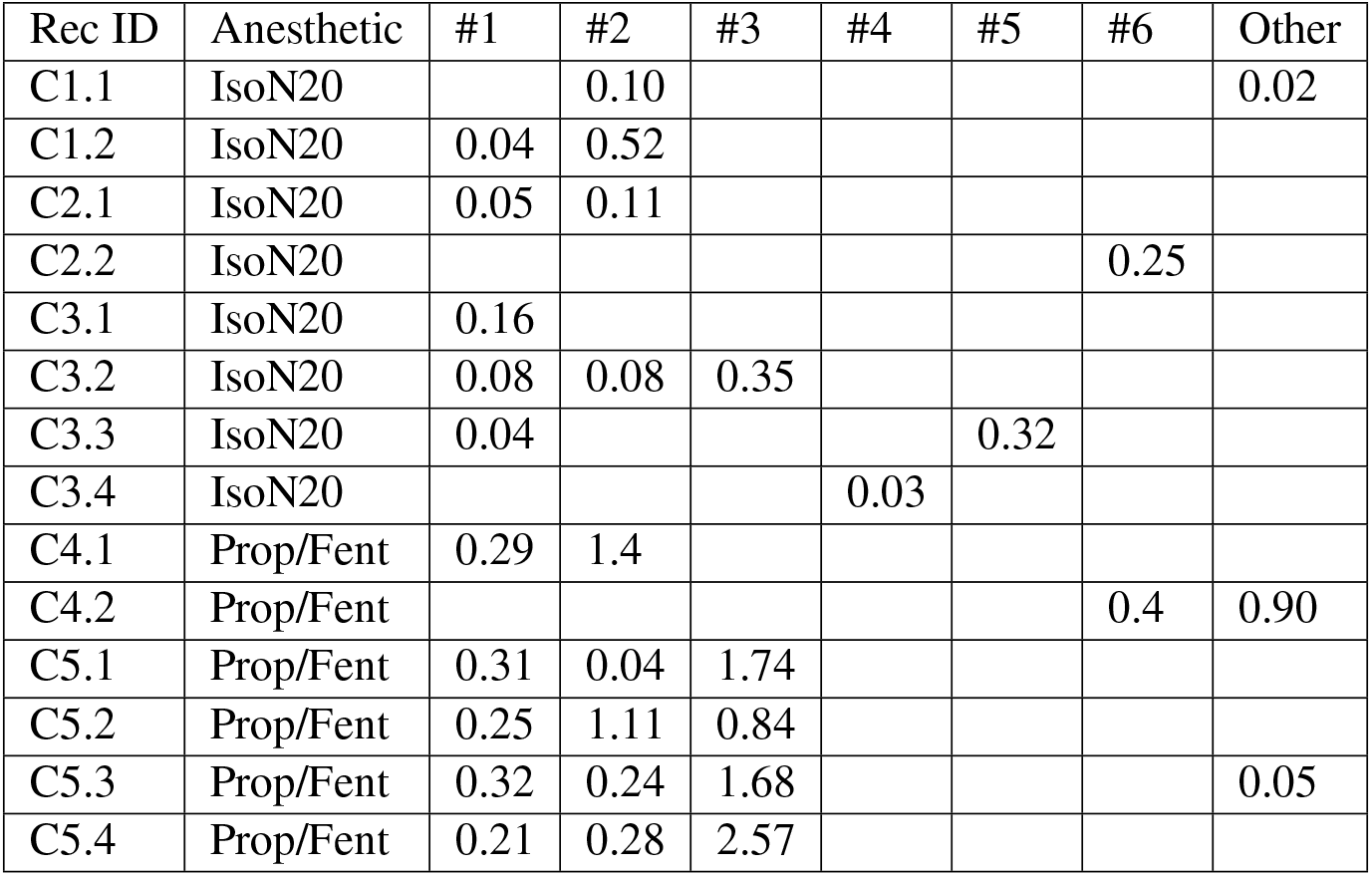
Cat V1 – LEC Firing Rates (Hz)

**Table 7:** Mouse Cortex – LEC Firing Rates (Hz)

| Rec ID | Anesthetic | Area | #1 | #2 | #3 | #4 | #5 | #6 | #7 | #8 |
| --- | --- | --- | --- | --- | --- | --- | --- | --- | --- | --- |
| MV1 | Isoflurane | Visual | 0.07 | 0.09 |  |  |  |  |  |  |
| MV2 | Isoflurane | Visual | 0.07 |  |  |  |  |  |  |  |
| MV3 | Isoflurane | Visual | 0.09 |  | 0.32 | 0.05 |  |  |  |  |
| MV4 | Isoflurane | Visual | 0.10 |  |  | 0.01 |  |  |  |  |
| MA1 | Isoflurane | Auditory |  |  |  |  | 0.06 | 0.01 |  |  |
| MA2 | Isoflurane | Auditory |  |  |  |  |  | 0.09 |  |  |
| MA3 | Isoflurane | Auditory |  |  |  |  |  |  | 0.37 |  |
| MB1 | Isoflurane | Barrel |  |  |  |  |  |  |  | 0.03 |
| Totals | 8 |  | 4 | 1 | 1 | 2 | 1 | 2 | 1 | 1 |

### LEC Events Can be Localised with 1 - 4ms Temporal Precision

We estimated the accuracy with which the times of individual LEC events could be measured by taking individual LEC templates and creating synthetic events by adding artificial noise with the same temporal structure as the real noise measured from the individual, aligned, events in the LEC (Fig 5). We then measured the change in the time of the event required to minimise the r.m.s. difference between it and the template, this change being the alignment error resulting solely from the added noise. The standard deviation of the resulting changes with repeated noise samples was taken as an estimate of the likely alignment error present with the real events in the sample. This was done for a number of different LECs in several different recordings. Error estimates ranged from 1.0 ms to 4.7 ms with a mean of 2.7 ms (n=11 in 6 different recordings). These estimates also put a lower bound on the accuracy with which spike times relative to individual LEC events can be measured.

**Figure 5.**
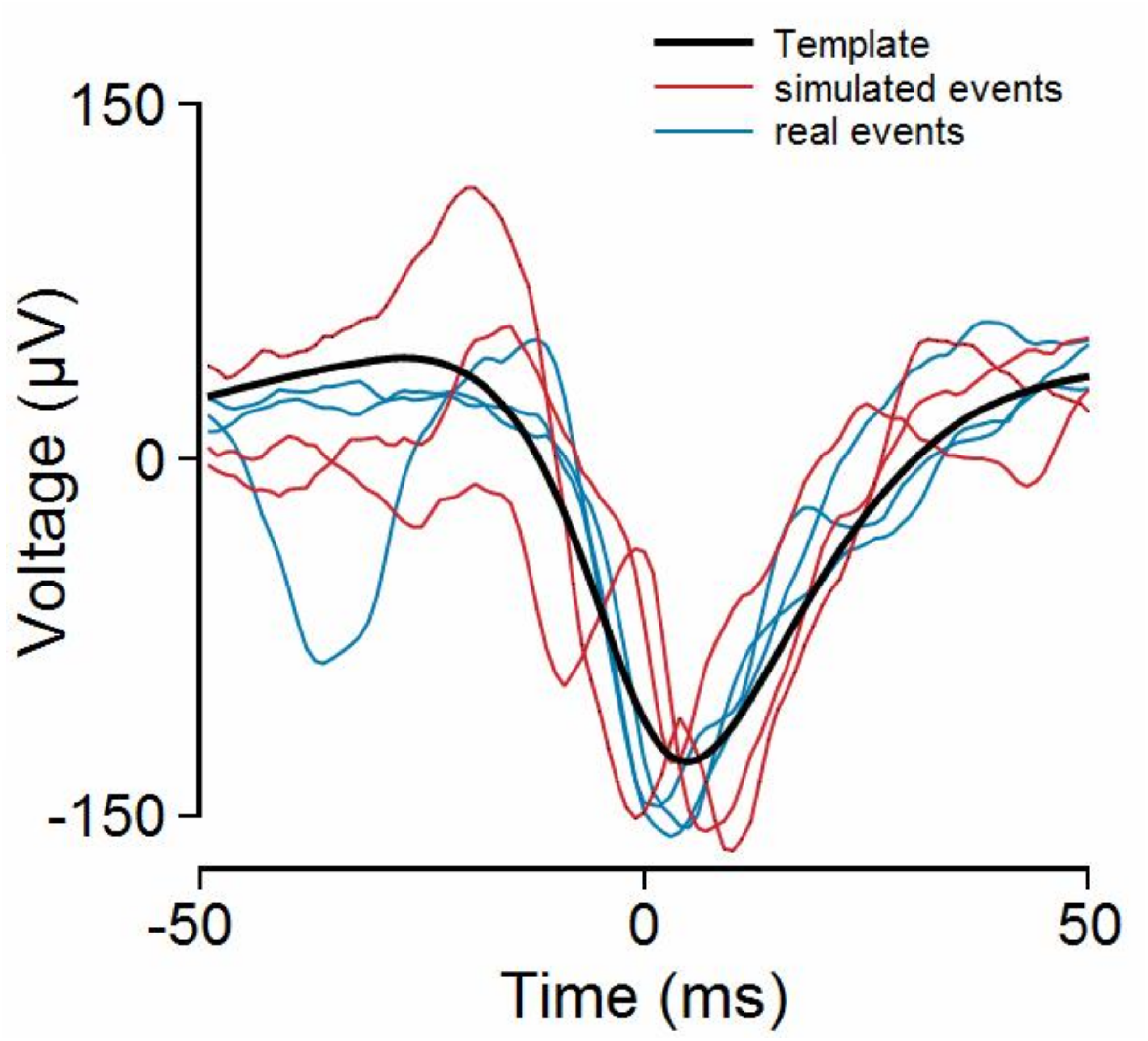
Estimating the temporal precision of individual LEC events. The solid black line shows the average (the template) of the events in a LEC. The 3 blue traces show the profiles of 3 individual events and the 3 red traces show the profiles of simulated events obtained by adding correlated noise to the template. Event times are estimated by finding the horizontal position of a profile that minimises the r.m.s. difference between it and the template. Time zero is defined as a weighted measure of the center of gravity of the template (see Methods).

The estimates were based on waveform data from single LFP channels (the one on which the peak-to-peak amplitude of the LEC template was a maximum). Adding additional channels improves the accuracy (though the improvements were found to be small, likely because signals tended to be highly correlated across adjacent channels). We also explored the temporal precision of LEC event detection by computing the stability of the full-width-half-max (FWHM) of the largest negative peak of each LEC event (Supplementary Figure 3). We found that the standard deviation of the FWHM were less than 10ms in most cases further suggesting that the overall shape of the LECs is relatively stable across time.

### Most Single Neurons Fire Precisely in Relation to LECs

We next considered the timing relation between single-unit firing and LEC events. We computed Peri-LEC-Event-Time-Histograms (PLETHs) using 5 ms bins and triggering off the LEC event times (Figure 6; see also methods). We found that neurons had qualitatively different PLETH distributions with different peaks shifted in time (Fig 6A: cat visual cortex recording; Fig 6B mouse visual cortex recording). Plotting the PLETHs for each neuron in a recording and arranging neurons by depth (Fig 6C-E: left) revealed a gradual temporal difference in peaks across almost all units recorded. Plotting units by depth (Fig 6C-E: right panels) revealed a preference for deeper units to fire earlier in the recording. We further analyzed whether deeper layer units fired before upper layer units, as reported by others (Sanchez-Vives and McCormick, 2000; Volgushev et al., 2006; Chauvette et al., 2010; Beltramo et al., 2013). PLETHs were ranked in order of depth (position of the unit) along the electrode (Figure 6C-E, rightmost panels). Units in the recordings were next divided into upper and lower halves (0 – 425 and 425 – 850 μm respectively for the mouse; and 0 – 750 and 750 – 1500 μm respectively for the cat recordings). We then counted the number of times the mean latency to the peak of the fitted gaussian was shorter for units in the upper vs. the lower halves of the recording (Supplementary Figure 5). In most cases latencies in the lower halves were shorter (10 of 14 mouse recordings and 21 of 32 cat recordings). These differences were not significant when tested separately (p = 0.09 and 0.06 respectively, binomial test) but were significant when both data sets were combined (p = 0.013, binomial test).

**Figure 6.**
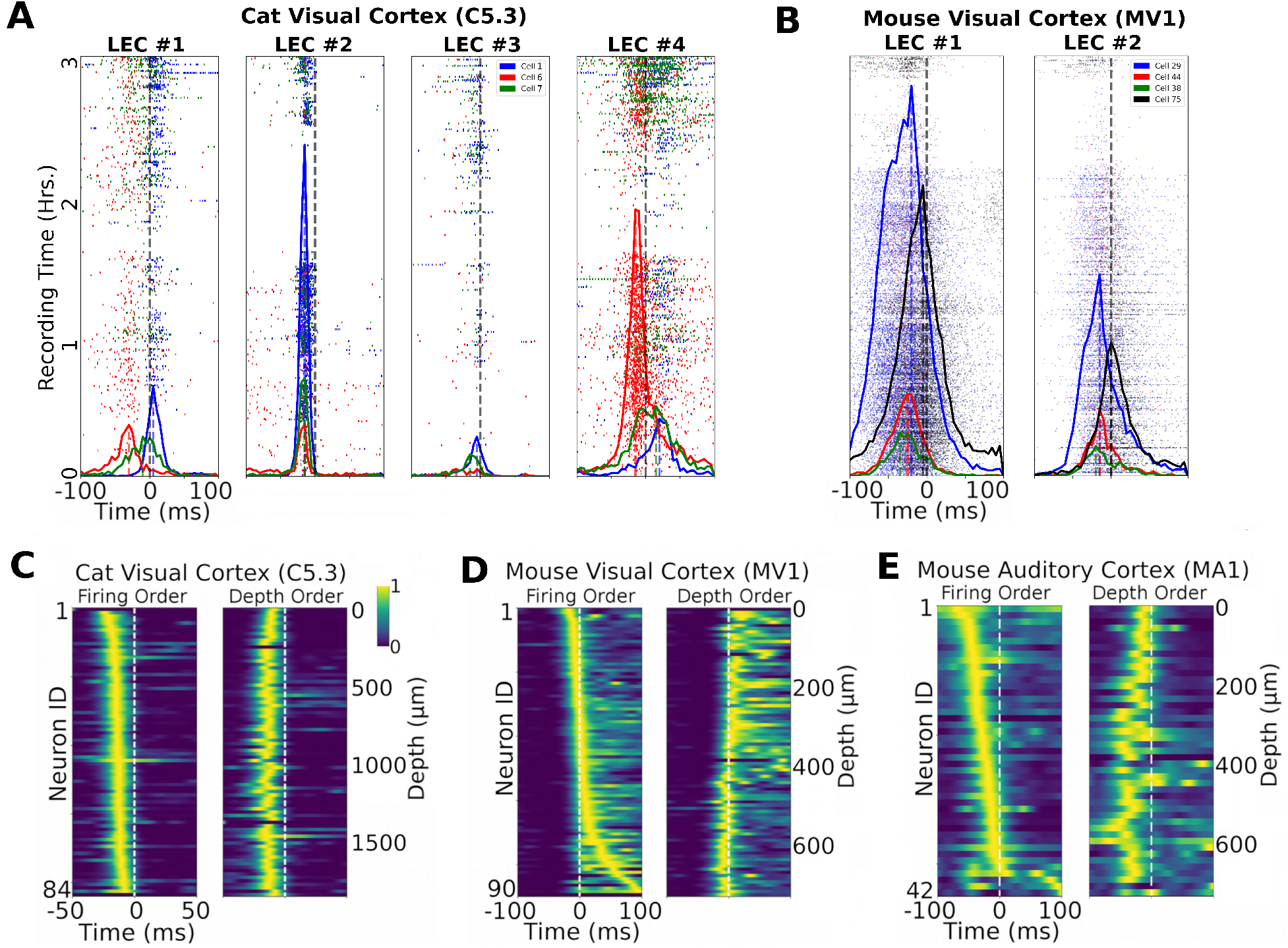
LECs Correlate with Single Neuron Activity. A. Peri-LEC event time histograms (PLETHs) for 3 example neurons and 4 LECs from a cat visual cortex recording (obtained as described in the Methods). Each dot represents a spike for a particular neuron (color) relative to the LEC event time (t=0 ms). B. Same as A but from a recording in mouse visual cortex. C. PLETHs for all neurons in A ordered by latency of the peak of each histogram relative to the time (t = 0) of each LEC-event (left; shortest latencies at the top) or by the depth of recorded neuron (right). D. Same as C but from a mouse visual cortex recording (MV1). E. Same as (C, D) but for a mouse auditory cortex recording (MA1)

We lastly considered whether the spiking distributions (i.e. not just the peaks) were statistically different across neurons. We found that despite sparse firing for many neurons, spiking distributions were almost always different between pairs of units recorded simultaneously (i.e. more than 90% of neuron pairs had < 0.01 pvalues; 2-sample Kolmogorov-Smirnov tests, Bonferroni corrected, Supplementary Figure 4). In other words, almost all neurons had unique firing distributions relative all other neurons even though their spiking distributions fell within a window of ~25-50 ms.

### Most Neurons Lock to LEC Defined UP-states with ±5-15 ms Latencies

In order to quantify the temporal precision with which different units could fire in relation to the LEC we selected units based on how well a gaussian fitted the PLETH (see Methods). Unit-PLETH pairs where the standard deviation of the fitted gaussian was > 50 ms showed poor fits and were excluded from analysis, as were units where no histogram bin exceeded a count of 5 in any 5 ms wide window (i.e. when only a few spikes occured during the recording period). These criteria resulted in the removal of ~25% of recorded units). We found most neurons had gaussian fits with 1-5ms standard deviations for both mouse (e.g. Fig 7A) and cat cortex (Fig 7B). Plotting the distribution of fits for all units recorded in different tracks and animals yielded an estimate of how ‘precise’ the firing of a unit could be determined relative the LEC. We found that across all unit-LEC pairs (3009) the mean precision was ±11.9 ms - though many units had lower values, with the lowest being 1.4 ms - close to the limit of the accuracy with which LEC event times could be measured.

**Figure 7.**
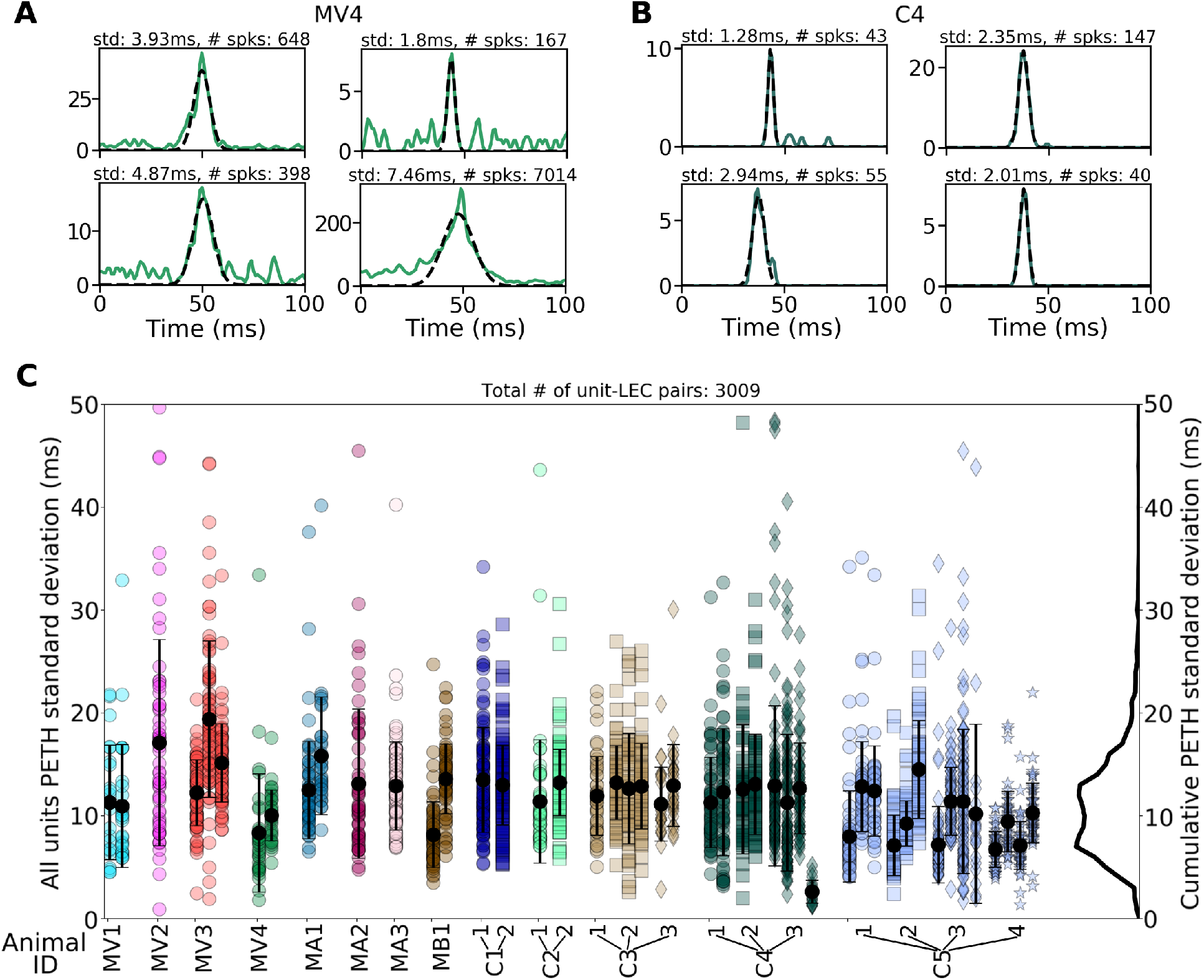
Single units fire precisely in relation to LEC onset. A. PLETH firing rate histograms (green lines) from a mouse visual cortex recording (ID: MV4) were fit with a Gaussian function (dashed black lines) to determine the mean latency and the width of the distribution. Fits were confined to units which fired reliably in relation to the events (see Results for details). B. Same as A, but for a cat visual cortex recording (ID: C4). C. All gaussian fits (3009) for each neuron-LEC pair in every track for all animal recordings. The vertical axis shows the width (sigma) of the fitted peak for each of the selected units. Each vertical column of points is the data from a single LEC type. Columns are grouped horizontally by animal and then by track number in each animal. The narrowest widths (indicating the most precisely firing units) are below 10 ms. The mean and mode of the overall distribution were 11.9 and 11.0 ms respectively.

### LECs Have Broad Mesoscale Correlates in Mouse Dorsal Cortex

We lastly sought to capture the meso-scale correlates of LECs by simultaneously recording widefield voltage-sensitive-dye (VSD) signals in mouse visual and auditory cortex (Fig 8A) and in GCaMP6s mice while recording extracellular potentials in mouse visual, barrel and auditory cortex (Fig 8B). LEC-triggered averages of widefield activity were computed as previously described (Xiao et al., 2017) for periods of 2 s relative to the LEC event time (i.e. ±2 s relative to each LEC event). In VSD recordings, the spatio-temporal patterns (termed ‘motifs’ here) showed a peak at the electrode recording site and revealed gradual multi-area cortical activation preceding LEC events (Fig 8A, t=0 s). This indicates that LECs are preceded by gradual membrane depolarizations and/or firing of many neurons as VSD activity represents both subthreshold and suprathreshold neural activity. This suggests that clustered LFP events are consistent with UP-state transition dynamics observed in intracellular recordings where near-simultaneous (10 - 100 ms) activation of neurons is observed during UP-state transitions across many cortical areas (Destexhe et al., 1999; Amzica and Steriade, 1995). The findings also suggest that LECs are the LFP-correlates of UP-state transitions in cortex. GCaMP6s mouse recordings also revealed a similar structure during LEC events (Figs 8B; see also Supplementary Videos 1 and 2).

**Figure 8.**
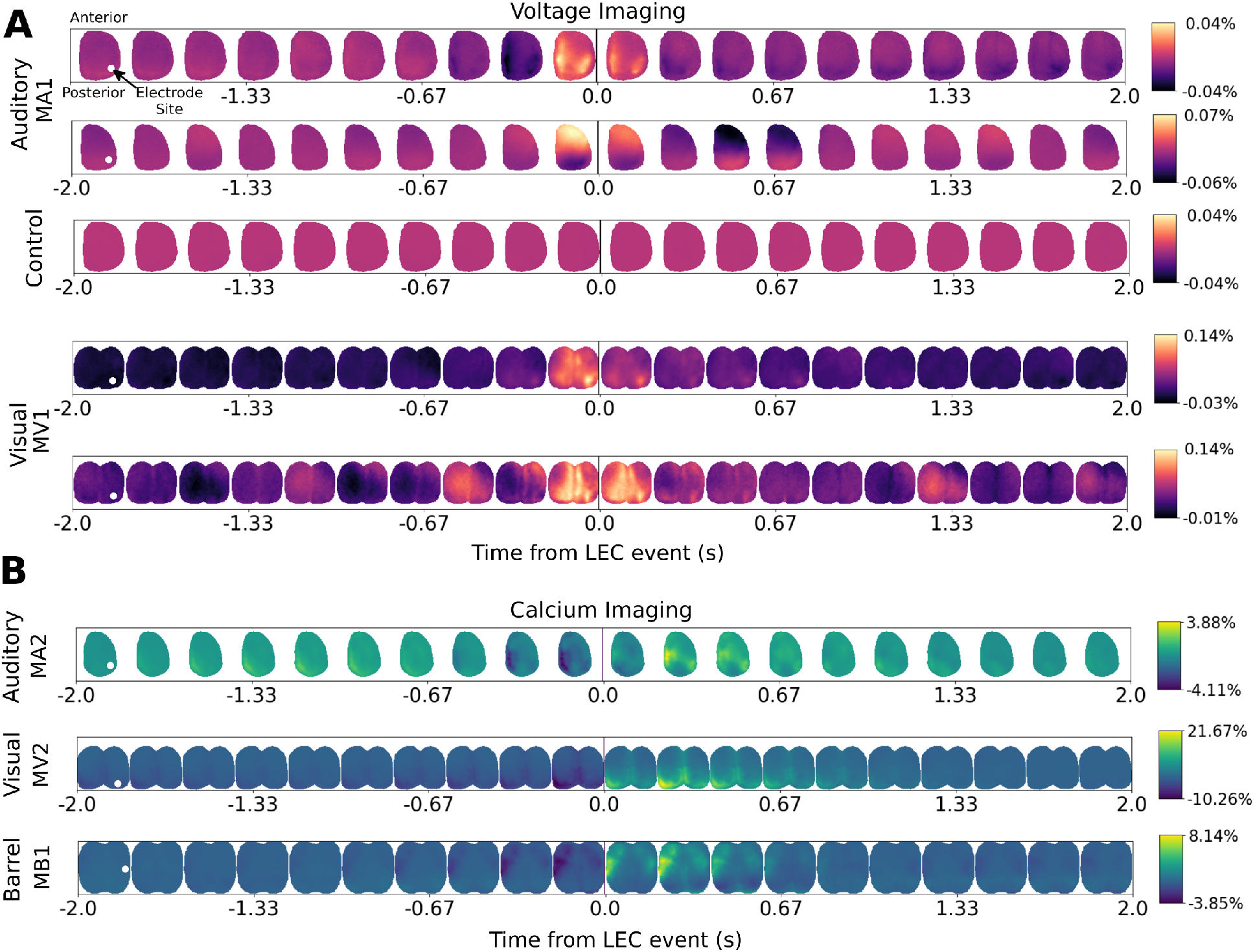
LEC-Triggered Single- and Bi-hemispheric Wide-field VSD Motifs. A. Voltage-sensitive-dye imaging of LEC-triggered dynamics in an auditory cortex recording reveals dynamics surrounding LEC event time for two different LECs. The control panel shows imaging data from randomly generated trigger times. Controls for other experiments were similar. B: Same as A but from two different recordings in mouse visual cortex. C: A, B but from GCaMP6s mice recordings in auditory, visual and barrel cortex (see also main text, Methods). White dots show the position of the recording electrode on which the triggering LECs were detected.

## Discussion

### Multiple LEC Types Suggests Multiple Sources of UP-state Genesis

It has been previously shown that UP-state transitions in single neurons have LFP correlates and various recent studies have used peaks in multi-unit-activity as markers of UP-state transitions (Amzica and Steriade, 1998; Chauvette et al., 2010; Saleem et al., 2010; Luczak et al., 2007). We describe a method for grouping UP-state transitions based on stereotyped multi-channel LFP event shapes. We show that in some recordings multiple classes of such events can be defined and that most single neurons synchronize their firing to within a few milliseconds of such events. The finding of multiple classes (1 to 4) of LECs in different species and cortical areas is consistent with a growing body of work identifying stereotypy in LFP recordings (Reichinnek et al., 2010; Ramirez-Villegas et al., 2015). Another study that used MUA to define UP-state transitions found two types of transitions in ketamine/xylazine-anesthetized rats based on UP-state duration (Luczak and Bartho, 2012). The findings of multiple classes of UP-state transitions is also supported by CSD correlates which have clearly different laminar patterns that are common across cortical areas and individuals of the same species. These findings of different patterns potentially occurring during UP-state transitions lend some support to the three cardinal oscillator hypothesis that UP-states can be caused and sustained by potentially independent cortico-thalamic-cortico populations (Crunelli and Hughes, 2010).

### Multi-channel LECs Capture Global UP-state Transitions

The term UP-state has commonly referred to a single neuron’s resting membrane potential transitioning from a hyperpolarized (i.e. non-spiking, e.g. −80 mV) to a depolarized (e.g. −60 mV) state (Steriade et al., 1993a). The term UP-state has also been used to refer to a global correlate of single neuron UP-state transitions where many (or possibly all) cortical neurons in a region depolarize and spike simultaneously (Neske, 2016). Defining an exact UP-state transition time faces certain problems. While most (or all) neurons depolarize simultaneously during an UP-state transition, not all neurons spike on every UP-state cycle (Volgushev et al., 2006; Chauvette et al., 2010). Additionally, individual neurons depolarize at different times and rates based on measurements of their intracellular membrane potentials (Lampl et al., 1999; Petersen et al., 2003) and can have different UP-state dynamics (Ros et al., 2009).

These factors make it challenging to track UP-states globally solely by recording a few neurons intracellularly or tracking peaks in MUA. In the present study we have shown that stereotypy in the LFP waveform can be used to establish the time of each event with a precision of a few milliseconds. This marker can then serve as a physiologically relevant temporal reference for evaluating single unit spike timing. Previous methods of identifying the time of UP-state transitions include measures based on changes in the firing rate of simultaneously recorded neurons (Luczak et al., 2007, 2009, 2013) but these methods have the limitation that they can only be applied to high-firing rate neurons (i.e. because peaks in cumulative firing rate histograms have low-SNR and are unreliable) in cortical areas that are not sparsely firing. Combined intracellular and LFP recordings (Chauvette et al., 2010) can be used to define UP-state transitions by fitting a sigmoid to LFP traces and defining a transition point at 10% of the amplitude of the sigmoid. The limitation is that both intracellular and extracellular recordings are required and that a somewhat arbitrary point is chosen as the time of the UP-state transition. (Saleem et al., 2010) used a method based on the phase of the LFP at frequencies below 4 Hz combined with multi-unit activity and single neuron recordings. This also has the limitation that both LFP and multi-unit activity are needed to define the onset of UP-states. Overall, none of the previous work has demonstrated a particular degree of precision in defining the time of UP-state onset.

While we claim to have found temporally precise markers of UP-state transitions our LEC times reflect a choice in event feature location (e.g. LEC t=0 ms can be chosen at peak, trough or centre-of-gravity of LEC event). Yet this limitation is also present in defining a global UP-state transition time using intracellular membrane potentials given that individual neurons can transition to UP-states at different times (Lampl et al., 1999; Petersen et al., 2003).

### Relation between LECs and K-complexes

Like the LECs studied here, K-complexes are transient large amplitude events that occur in EEG or LFP recordings during the synchronised state in anesthesia and during stage 2 slow wave sleep (e.g. (Loomis et al., 1937; Amzica and Steriade, 1998). In humans, K-complexes typically last up to 1 second and occur every 1 – 2 minutes. They are often followed by sleep spindles - a burst of rapid oscillations at a frequency of 10 – 12 Hz. In recordings from ketamine-anesthetised rats (Luczak and Bartho, 2012) also describe large amplitude fluctuations in LFP recordings which mark transitions to UP-states where large numbers of neurons start firing at about the same time. That study suggests that these fluctuations are homologous to K-complexes. However there are differences between all three sets of observations – classical sleep related K-complexes in humans, those of (Luczak and Bartho, 2012) and ours. Like Luczak and Bartho’s findings, ours differ from classical K-complexes in being much faster (lasting 100 – 200 ms compared to up to 1 second) and occurring at much higher rates (several/second compared to every 1 – 2 minutes). Ours also differ from those observed by Luczak and Bartho in being simpler in structure, with typically a single large peak (positive or negative) flanked by two of opposite sign (see Fig 1G). We also did not find the traveling wave events described by Luczak and Bartho, possibly because these are reported to be of smaller amplitude and they perhaps fell below our detection threshold. Nor did we observe obvious sleep spindles following our events. Reasons for these various differences would include species (human K-complexes and spindles may be generally slower than in rats and cats), the fact that the animals were not naturally sleeping, types of anesthesia (Luczak and Bartho used ketamine, we used either isoflurane or propofol) and cortical area (we studied visual areas whereas Luczak and Bartho studied rat auditory cortex). A conservative hypothesis that might reconcile all of these findings is that K-complexes constitute a large and heterogeneous class of high amplitude transient activity in the LFP associated with UP-state transitions and widespread firing of neurons. Our findings of multiple types of LECs support such heterogeneity within single cortical areas and recording sessions, as well as suggesting that specific types of events may be identifiable within areas and across different individuals of the same species (Fig. 4).

### LECs as temporally precise global markers of UP-state transitions

As an alternative global-definition of UP-state transitions, LECs have advantages over single neuron patch clamp recordings in that they can be more rigorously defined using statistical clustering methods while also being more stable as they consist of spatially broad (i.e. 100 μm to 1000 μm) LFP contributions from multiple sources (Buzsáki et al., 2012) while being largely independent of any single neuron’s activity. While simultaneous intracellular recordings from many neurons might eventually be feasible, defining global UP-state transitions using such recordings stills requires averaging UP-state transition times leading to a definition that is dependent on the particular set of recorded neurons. Since the LFP represents the activity of a large population of neurons, transition times estimated from the stereotyped shapes of multi-channel LFP signals may provide a principled and non-circular methodology, i.e. it does not define UP-state transition spiking based on the cumulative spiking of many neurons.

We propose that future work should focus on the implications of the narrowly defined UP-state transitions spiking (previous work showed histogram widths of 20-150 ms (Luczak et al., 2007)). Such narrower spiking distributions lend support to spike timing and firing order being present in cortical processing (Panzeri et al., 2001; Gautrais and Thorpe, 1998).

---

**Figure 1-1:**
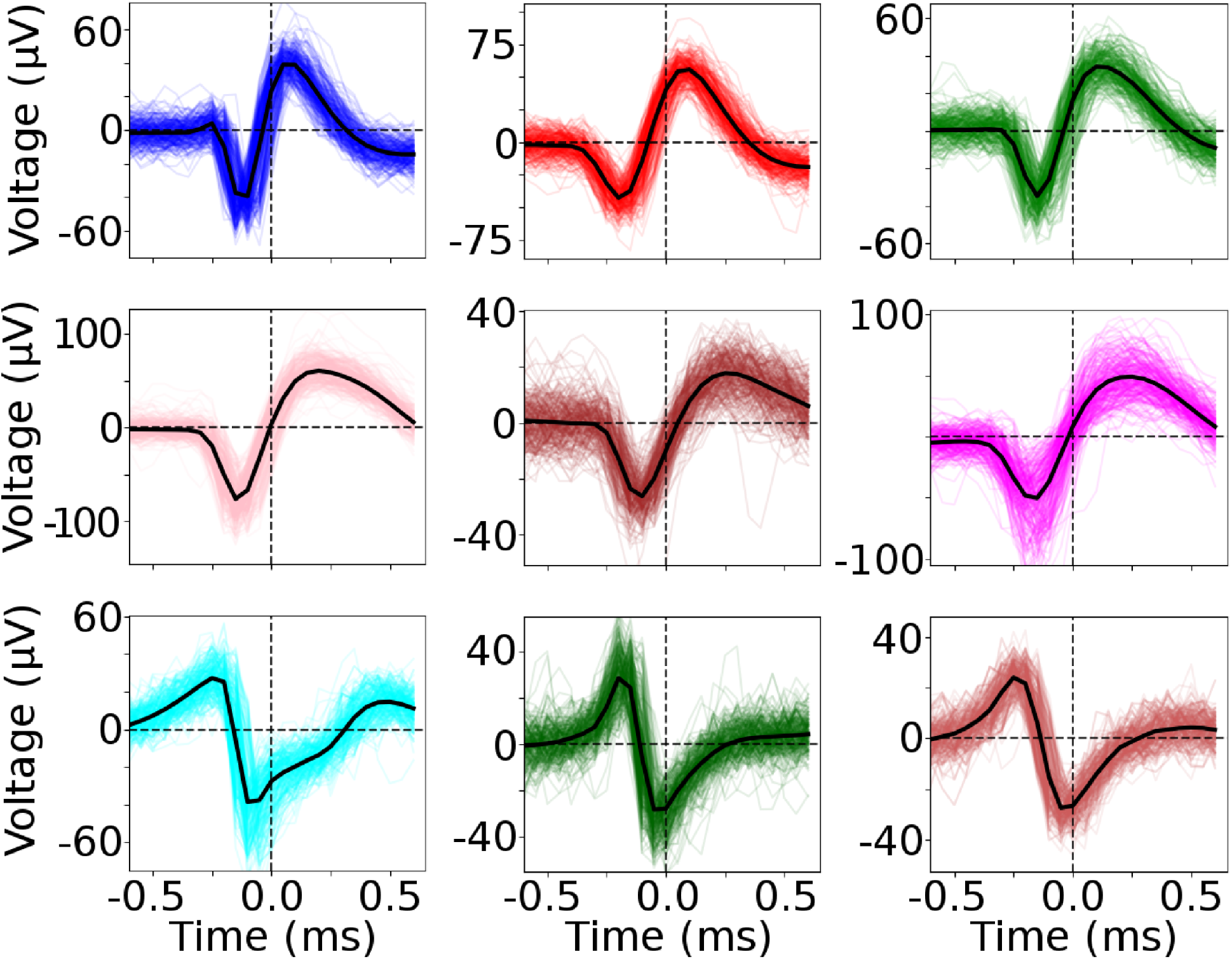
Examples of single unit clustered spikes. Spikes on the maximum amplitude channels for nine randomly chosen single units from both cat and mouse recordings.

**Figure 3-1:**
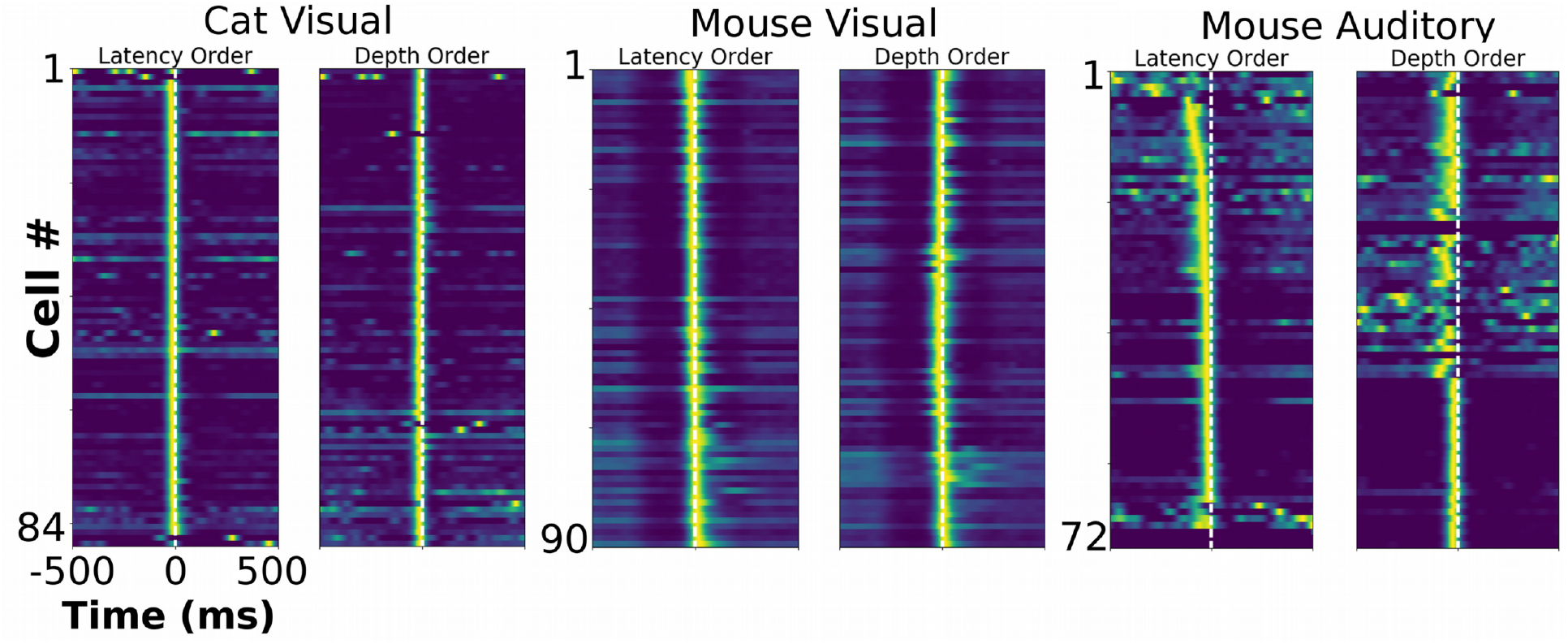
PETH distributions computed for wider temporal windows (−500 ms to +500 ms) than Figure 6, for 3 LECs recorded in cat visual cortex, mouse visual cortex and mouse auditory cortex.

**Figure 5-1:**
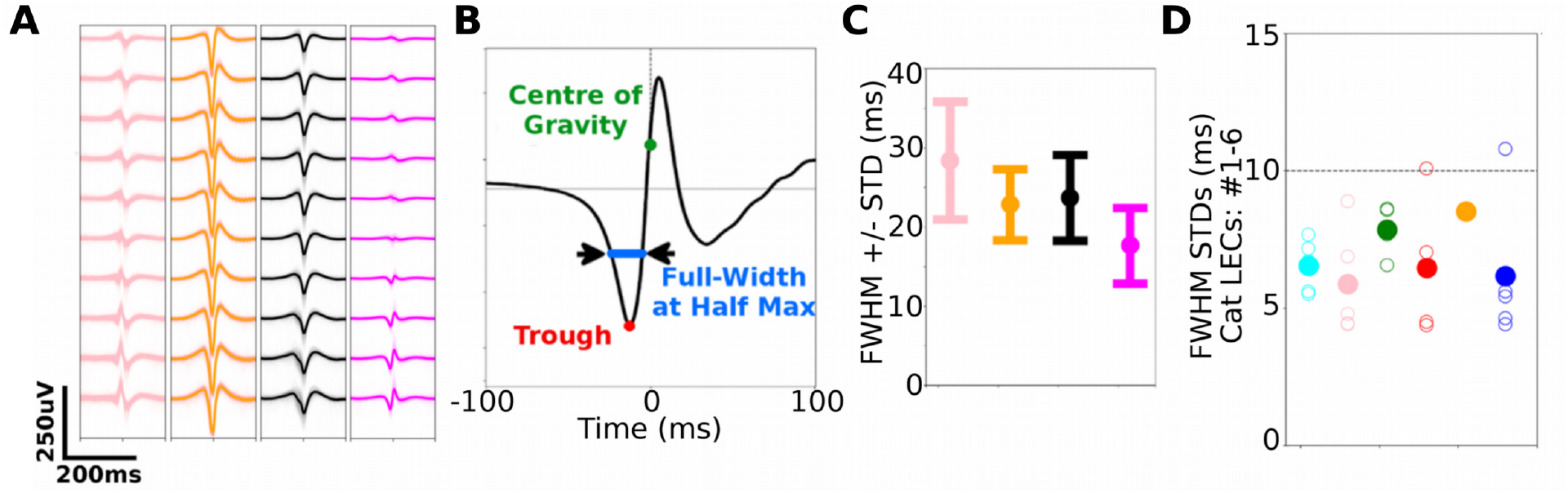
Measuring the stability of LEC events. A. The four LEC templates shown in Fig 1G. B. Measurement of the full-width-half-max (FWHM) from the first trough (i.e. negative peak) of each LEC event. C. FWHM means and standard deviations of the four LEC events shown in A reveal that most LEC’s FWHM standard deviations are <10 ms. D. Same as C but for the first 6 LEC groups in Fig 4, reveal the vast majority of LECs have individual standard deviations < 10 ms.

**Figure 6-1.**
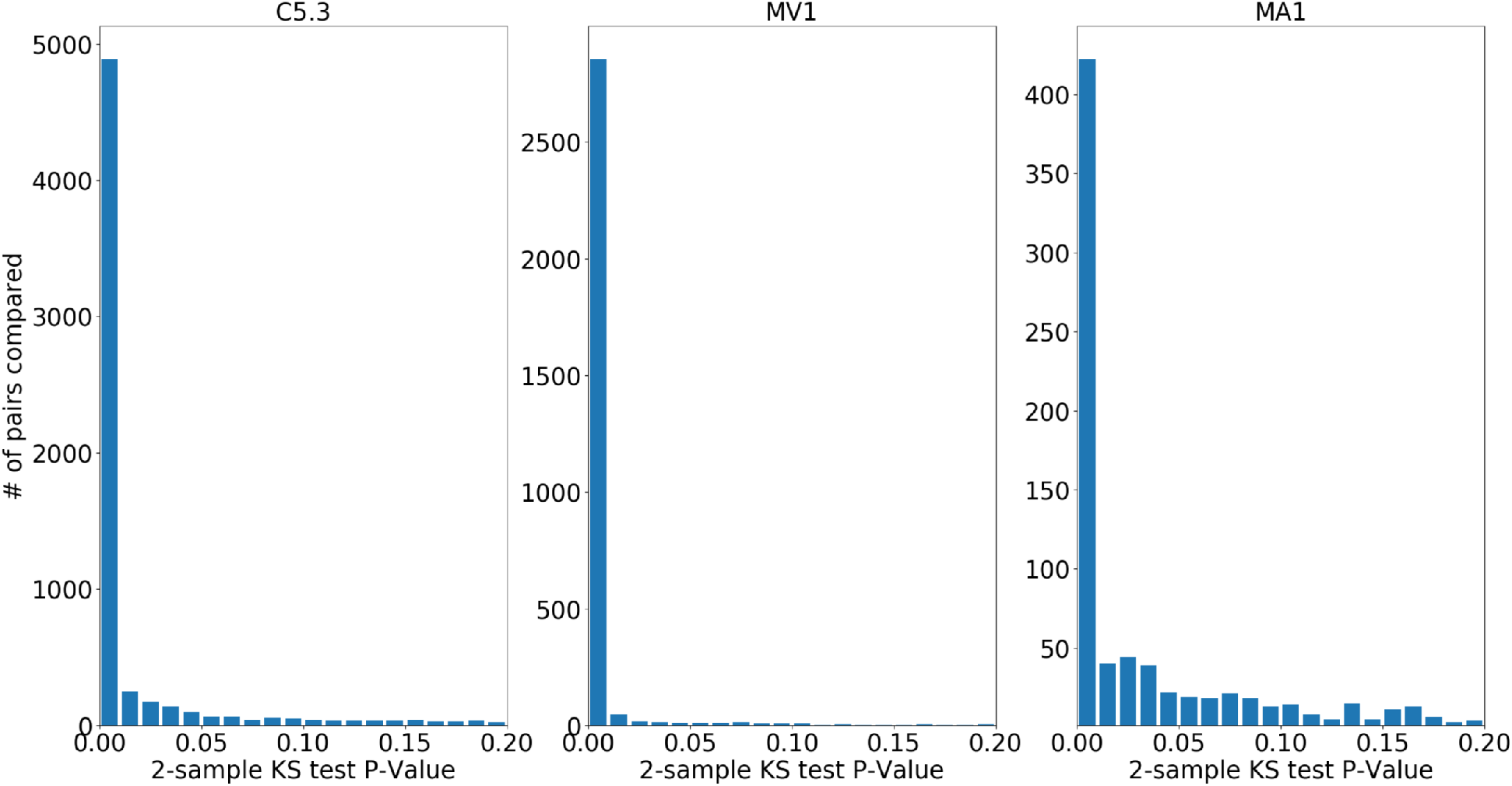
LEC-triggered spiking distributions are significantly different across neurons. The significance of the difference in the distributions between pairs of neurons was assessed using 2 sample Kolmogorov-Smirnov tests with Bonferonni correction. Histograms show the distributions of *p* values for all the unit pairs in particular recordings A: recording C5.3; B recording MV1 and C recording MA1.

**Figure 6-2.**
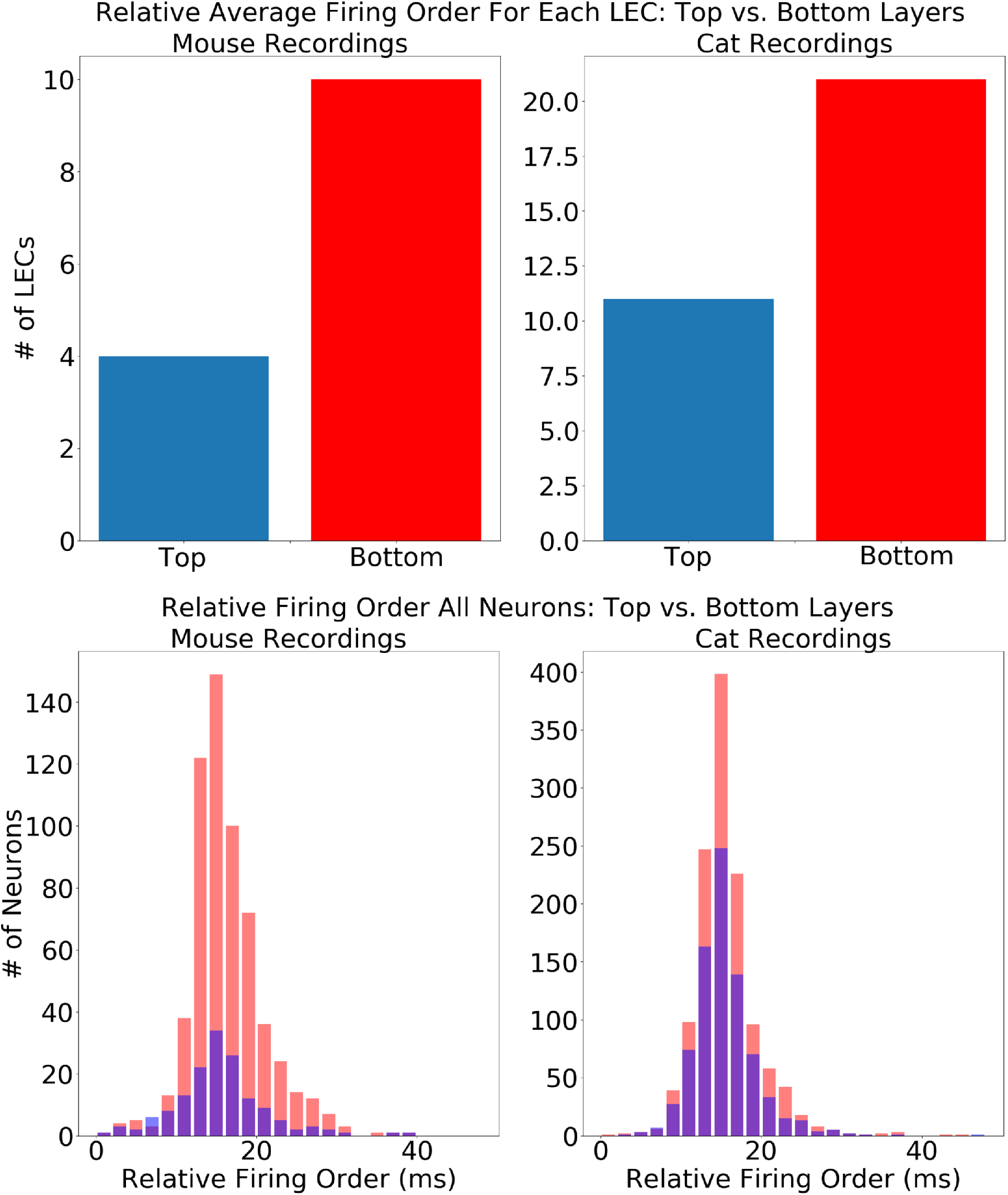
Deeper layer neurons are more likely to fire first during LEC-events. Top: histograms show the number of LECs in which superficial layer (blue) neurons (0-425 *μ*m: mouse; 0-750 *μ*m: cat) spiked before deeper layer (red) neurons (425-850 *μ*m: mouse; 750-1500 *μ*m: cat). Order was based on the means of Gaussian fits to LEC-triggered firing rate histograms. There is an overall bias for deeper layer neurons to spike first (see text for further description of statistical tests). Bottom: pooling all neuron relative firing times did not reveal substantial order differences between superficial layer neurons (blue) or deeper layer neurons (red) firing first. This was likely due to individual LECs eliciting different lag spiking which is averaged out when pooling all relative spiking times across all LECs.

## Acknowledgements

This work was supported by: Canadian Institutes of Health Research (CIHR) Operating Grant MOP-15360 and National Science and Engineering Research Council of Canada 178702 to N.V.S.; Canadian Institutes of Health Research (CIHR) Operating Grant MOP-12675 and Foundation Grant FDN-143209 to T.H.M. We thank Pumin Wang and Cindy Jiang for surgical assistance and Jamie Boyd for technical assistance.

## References

Amzica F, Steriade M (1995) Short- and long-range neuronal synchronization of the slow (¡ 1 hz) cortical oscillation. Journal of Neurophysiology 73:20–38.

Amzica F, Steriade M (1998) Electrophysiological correlates of sleep delta waves. Electroencephalogr Clin Neurophysiol 107:69–83.

Beltramo R, D’Urso G, DalMaschio M, Farisello P, Bovetti S, Clovis Y, Lassi G, Tucci V, DePietriTonelli D, Fellin T (2013) Tbd. Nature Neuroscience 16:227–234.

Bermudez-Contreras E, Schjetnan A, Muhammad A, Bartho P, McNaughton B, Kolb B, Gruber A, Luczak A (2013) Formation and reverberation of sequential neural activity patterns evoked by sensory stimulation are enhanced during cortical desynchronization. Neuron 79:555–566.

Buzsáki G, Anastassiou CA, Koch C (2012) The origin of extracellular fields and currents — EEG, ECoG, LFP and spikes. Nat Rev Neurosci 13:407–420.

Chauvette S, Volgushev M, Timofeev I (2010) Origin of active states in local neocortical networks during slow sleep oscillation. Cerebral Cortex 10:2660–2674.

Crunelli V, Hughes S (2010) The slow (<1 hz) rhythm of non-rem sleep: a dialogue between three cardinal oscillators. Nature Neuroscience 13:9–17.

Destexhe A, Contreras D, Steriade M (1999) Spatiotemporal analysis of local field potentials and unit discharges in cat cerebral cortex during natural wake and sleep states. J Neurosci 19:4595–4608.

Gautrais J, Thorpe S (1998) Rate coding versus temporal order coding: a theoretical approach. Biosystems 48:57–65.

Haider B, Duque A, Hasenstaub AR, McCormick DA (2006) Neocortical network activity in vivo is generated through a dynamic balance of excitation and inhibition. J Neurosci 26:4535–4545.

Harris KD, Thiele A (2011) Cortical state and attention. Nat Rev Neurosci 12:509–523.

Jun J, Mitelut C, Lai C, Gratiy S, Anastassiou C, Harris T (2017) Real-time spike sorting platform for high-density extracellular probes with ground-truth validation and drift correction. bioRxiv.

Lampl I, Reichova I, Ferster D (1999) Synchronous membrane potential fluctuations in neurons of the cat visual cortex. Neuron 22:361–374.

Li CY, Poo MM, Dan Y (2009) Burst spiking of a single cortical neuron modifies global brain state. Science 324:643–646.

Loomis A, Harvey E, Hobart G (1937) Cerebral states during sleep, as studied by human brain potentials. J Exp Psychol 21:127–44.

Luczak A, Bartho P, Harris KD (2013) Gating of sensory input by spontaneous cortical activity. J Neurosci 33:1684–1695.

Luczak A, Barthó P, Marguet SL, Buzsáki G, Harris KD (2007) Sequential structure of neocortical spontaneous activity in vivo. PNAS 104:347.

Luczak A, McNaughton B, Harris K (2015) Packet-based communication in the cortex. Nature Reviews Neuroscience 16:745–755.

Luczak A, Bartho P, Harris K (2009) Spontaneous events outline the realm of possible sensory responses in neocortical populations. Neuron 413-425:413–425.

Luczak A, Bartho P (2012) Consistent sequential activity across diverse forms of up states under ketamine anesthesia. European Journal of Neuroscience 36:2830–2838.

McCormick D, Shu Y, Hasenstabu A (2004) Balanced recurrent excitation and inhibition in local cortical networks. Excitatory-Inhibitory Balance: Synapses, Circuits, Systems pp. 113–122.

McCormick D, Yuste R (2006) Up states and cortical dynamics. Microcircuits: the Interface between Neurons and Global Brain Function pp. 327–346.

Mohajerani M, Chan A, Mohsenvand M, LeDue L, Liu R, McVea D, Boyd J, Wang Y, Murphy T, Reimers M (2013) Spontaneous cortical activity alternates between motifs defined by regional axonal projections. Nature Neuroscience 16:10.

Mohajerani M, McVea D, Fingas M, Murphy T (2010) Mirrored bilateral slow-wave cortical activity within local circuits revealed by fast bihemispheric voltage-sensitive dye imaging in anesthetized and awake mice. Journal of Neuroscience 30:3745–3751.

Neske GT (2016) The slow oscillation in cortical and thalamic networks: Mechanisms and functions. Frontiers in Neural Circuits 9:88.

Nicholson C, Freeman J (1975) Theory of current source-density analysis and determination of conductivity tensor for anuran cerebellum. Journal of Neurophysiology 38(2):356–68.

Pachitariu M, Stringer C, Dipoppa M, Schröder S, Rossi LF, Dalgleish H, Carandini M, Harris KD (2017) Suite2p: beyond 10,000 neurons with standard two-photon microscopy. bioRxiv.

Panzeri S, Petersen RS, Schultz SR, Lebedev M, Diamond ME (2001) The role of spike timing in the coding of stimulus location in rat somatosensory cortex. Neuron 29:769–777.

Petersen CCH, Hahn TTG, Mehta M, Grinvald A, Sakmann B (2003) Interaction of sensory responses with spontaneous depolarization in layer 2/3 barrel cortex. PNAS 100:13638–13643.

Quian Quiroga R, Nadasdy Z, Ben-Shaul Y (2004) Unsupervised spike detection and sorting with wavelets and superparamagnetic clustering. Neural Comput 16:1661–1687.

Ramirez-Villegas JF, Logothetis NK, Besserve M (2015) Diversity of sharp-wave–ripple lfp signatures reveals differentiated brain-wide dynamical events. Proceedings of the National Academy of Sciences 112:E6379–E6387.

Reichinnek S, Künsting T, Draguhn A, Both M (2010) Field potential signature of distinct multicellular activity patterns in the mouse hippocampus. Journal of Neuroscience 30:15441–15449.

Ros H, Sachdev RNS, Yu Y, Šestan N, McCormick DA (2009) Neocortical networks entrain neuronal circuits in cerebellar cortex. Journal of Neuroscience 29:10309–10320.

Saleem AB, Chadderton P, Apergis-Schoute J, Harris KD, Schultz SR (2010) Methods for predicting cortical UP and DOWN states from the phase of deep layer local field potentials. J Comput Neurosci 29:49–62.

Sanchez-Vives M, Massimini M, Mattia M (2017) Shaping the default activity pattern of the cortical network. Neuron 94:993–1001.

Sanchez-Vives M, McCormick D (2000) Cellular and network mechanisms of rhythmic recurrent activity in neocortex. Nature Neuroscience 3:1027–1034.

Shoham D, Glaser DE, Arieli A, Kenet T, Wijnbergen C, Toledo Y, Hildesheim R, Grinvald A (1999) Imaging cortical dynamics at high spatial and temporal resolution with novel blue voltage-sensitive dyes. Neuron 24:791–802.

Sirota A, Buzsaki G (2005) Interaction between neocortical and hippocampal networks via slow oscillations. Thalamus and related systems 3:245–259.

Sirota A, Csicsvari J, Buhl D, Buzsáki G (2003) Communication between neocortex and hippocampus during sleep in rodents. Proceedings of the National Academy of Sciences 100:2065–2069.

Steriade M (2001) Impact of network activities on neuronal properties in corticothalamic systems. Journal of Neurophysiology 86:1–39.

Steriade M, Nuñez A, Amzica F (1993a) A novel slow (<1 Hz) oscillation of neocortical neurons in vivo: depolarizing and hyperpolarizing components. J Neurosci 13:3252–3265.

Swindale NV, Spacek MA (2015) Spike detection methods for polytrodes and high density microelectrode arrays. J Comput Neurosci 38:249–261.

Swindale N, Spacek M (2014) Spike sorting for polytrodes: a divide and conquer approach. Frontiers in Systems Neuroscience 8:6.

Vanni M, Murphy T (2014) Mesoscale transcranial spontaneous activity mapping in gcamp3 transgenic mice reveals extensive reciprocal connections between areas of somatomotor cortex. Journal of Neuroscience 34:15931–15946.

Volgushev M, Chauvette S, Mukovski M, Timofeev I (2006) Precise long-range synchronization of activity and silence in neocortical neurons during slow-wave sleep. Journal of Neuroscience 26:5665–5672.

Wilson FA, O’Scalaidhe SP, Goldman-Rakic PS (1994) Functional synergism between putative *γ*-aminobutyrate-containing neurons and pyramidal neurons in prefrontal cortex. PNAS 91:4009–4013.

Xiao D, Vanni M, Mitelut C, Chan A, LeDue J, Xie Y, Chen A, Swindale N, Murphy T (2017) Mapping cortical mesoscopic networks of single spiking cortical or sub-cortical neurons. eLife 6:e19976.

